# Systematic integration of protein affecting mutations, gene fusions, and copy number alterations into a comprehensive somatic mutational profile

**DOI:** 10.1101/2022.07.22.501139

**Authors:** Shawn S Striker, Sierra F Wilferd, Erika M Lewis, Samantha A O’Connor, Christopher L Plaisier

## Abstract

Somatic mutations occur as random genetic changes in genes through protein-affecting mutations (PAMs), gene fusions, or copy number alterations (CNAs). Mutations of different types can have a similar phenotypic effect (i.e., allelic heterogeneity) and should be integrated into a unified gene mutation profile. We developed OncoMerge to fill this niche of integrating somatic mutations to capture allelic heterogeneity, assign a function to mutations, and overcome known obstacles in cancer genetics. Application of OncoMerge to the TCGA Pan-Cancer Atlas increased the number and frequency of somatically mutated genes and improved the prediction of the somatic mutation role as either activating or loss of function. Using integrated somatic mutation matrices increased the power to infer gene regulatory networks and uncovered the enrichment of switch-like feedback motifs and delay-inducing feed-forward loops. These studies demonstrate that OncoMerge efficiently integrates PAMs, fusions, and CNAs and strengthens downstream analyses linking somatic mutations to cancer phenotypes.

**Motivation:** Genes are mutated in tumors through either PAMs, CNAs, or gene fusions, thereby splitting the signal of the somatic mutation effect across these three mutation types. We developed OncoMerge to systematically integrate the three mutation types into a single mutation profile that better captures the impact of somatic mutations on cancer phenotypes. As a tool OncoMerge fills the gap between the sophisticated variant calling pipelines and downstream analyses.

## Introduction

The accumulation of somatic mutations in patient tumors drives and reinforces cancer phenotypes. The three main types of somatic mutations that modify the function of a gene or render it non-functional are: 1) protein affecting mutations (PAMs), 2) gene fusions, and 3) copy number alterations (CNAs). A PAM is a point mutation, short insertion, or short deletion inside a gene’s coding region or splice sites^1^. Gene fusions occur when genomic rearrangements join two genes into a novel chimeric gene or place a promoter in front of a new gene, causing misexpression^2^. Finally, CNAs occur frequently in tumors where whole chromosomes, chromosomal arms, or localized genomic segments are duplicated or deleted^3, 4^. Somatic mutation via PAM, gene fusion, or CNA can have similar effects on cancer phenotypes, i.e., allelic heterogeneity. This interchangeability and the erratic circumstances that produce somatic mutations lead to the mixture of mutation types observed in large cohorts of patient tumors^1^.

Describing how somatic mutations in a gene impact cancer phenotypes requires integrating the information from all three mutation types. Most studies linking somatic mutations to cancer phenotypes focus on one mutation type. This leads to missing associations for mutations primarily found in another type and reduced power to detect associations for mutations with high allelic heterogeneity that span the mutation types. Thus, a current obstacle facing those studying the downstream effects of somatic mutations is the lack of an established method for integrating PAMs, gene fusions, and CNAs into a comprehensive gene mutation profile. The lack of integration methods is due to several complicating factors. Firstly, the allelic heterogeneity observed in and between tumors means that different mutations in the same gene can be equivalently oncogenic. Second, it is challenging to discern driver (causal) from passenger (non-causal) somatic mutations. Third, an algorithm must be able to systematically integrate the binary PAM and gene fusion (mutated or not) with the quantitative copy number from CNAs. Lastly, some tumors have drastically higher somatic mutation rates than others (e.g., microsatellite instability^5^ and hypermutation^6^). These higher mutation rates confound any frequency-based integration approach and drive the discovery of spurious somatic mutations. We developed OncoMerge to fill the somatic mutation integration niche by providing an algorithm that systematically overcomes these obstacles to generate an integrated gene mutation profile. The input for the OncoMerge algorithm is the output from state-of-the-art methods for detecting PAMs (MC3^1^ and MutSig2CV^7^), transcript fusions (PRADA^2, 8^), and CNAs (GISTIC2.0^9^). Each method provides the likelihood that a somatic mutation happens by chance alone. Filtering on these statistics focuses integration efforts on genes most likely to harbor functional mutations. The integrated mutation profiles improve the power to detect associations with cancer phenotypes leading to a more comprehensive understanding of how genetic alterations drive cancer phenotypes.

The tremendous amount of cancer genome sequencing data generated in the last ten years has enabled efforts to discover and catalog somatic mutations across many cancers^1, 10^. Many algorithms have been developed to discern which somatic mutations are drivers, how the mutations affect genes^6, 7, 11–17^, and databases to search and view somatically mutated driver genes^16–18^. There also exist approaches for integrating somatic mutations. OncoPrint from cBioportal^18^ can visually overlay somatic mutation types across patient tumors for a gene of interest. The OncodriveROLE algorithm^13^ was developed to discover driver genes by systematically integrating PAMs and CNVs. However, neither OncoPrint nor OncodriveROLE provides an integrated mutational profile that can be used in downstream analyses. The impact of somatic mutations can be classified as activating (Act) gene function (typically found in oncogenes) or loss of function (LoF) (typically found in tumor suppressor genes)^13^. It has also been demonstrated that the systematic integration of PAM and CNA somatic mutations for a gene improves the ability to determine Act or LoF status^13^. These foundational studies have created a platform to develop an algorithm that systematically integrates the three somatic mutation types.

The systematic integration of somatic mutations requires choosing a gene-level model that determines how the data for the three somatic mutation types will be integrated, the somatic mutation role. We determine the somatic mutation role by employing rules similar to those in OncodriveROLE^13^ (**Figure 1**). The possible somatic mutation roles in OncoMerge are PAM, Fusion, CNA amplification (CNAamp), CNA deletion (CNAdel), Act, or LoF. The PAM, Fusion, CNAamp, and CNAdel somatic mutation roles use the unintegrated somatic mutation profile for the chosen role in the final mutation matrix. The Act and LoF are integrated mutation roles that harness allelic heterogeneity. Allelic heterogeneity is especially prevalent in tumor suppressor genes, where mutations at many positions in a gene disrupt its function to prevent cancer phenotypes^3^. Allelic heterogeneity is less prevalent for oncogenes where a small number of specific gain of function alleles are needed to drive cancer phenotypes^3^. Genes underlying CNAs can add another layer of information as tumor suppressors are often deleted, which has an equivalent oncogenic effect as missense or truncating PAMs. The LoF role is designated when PAMs, Fusions, and CNAdels are integrated. Oncogenes are often amplified as this typically leads to overexpression of the underlying genes, which has a similar positive effect on gene function as a gain of function PAM. The Act role is designated when PAMs, Fusions, and CNAamps are integrated. Systematic determination of the somatic gene role and application of the rules laid out above are used to integrate the three mutation types into a comprehensive somatic mutation profile.

**Figure 1.**
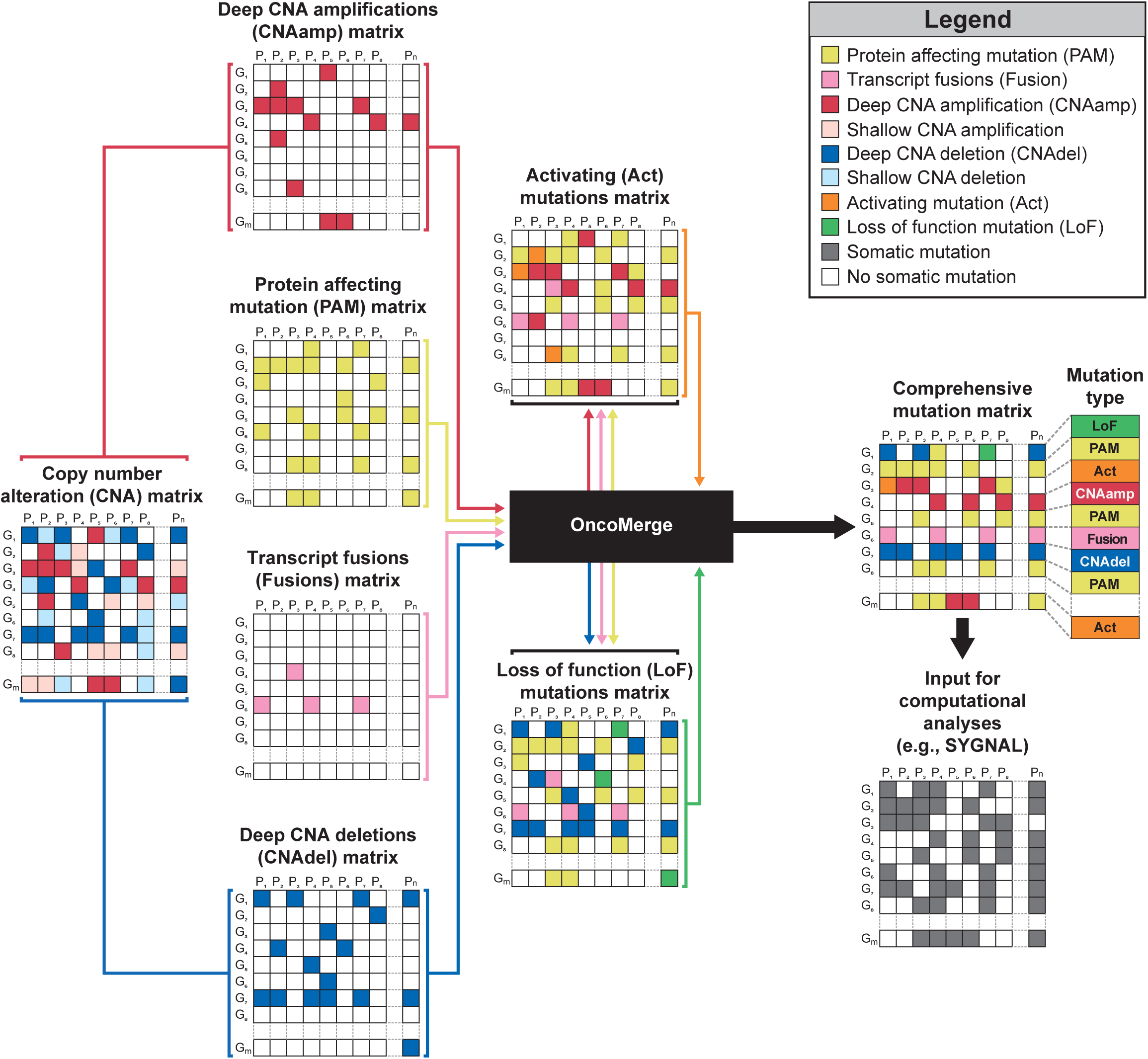
OncoMerge integrates PAMs, fusions, and CNAs into an integrated mutation matrix with the most suitable mutation type for each gene. The input data for OncoMerge includes the PAM, transcript fusion, and CNA matrices. OncoMerge then generates six matrices (PAM, Fusion, CNAamp, CNAdel, Act, and LoF) and uses mutational frequency and statistical filters to determine each gene’s most suitable somatic mutation role.

The lists of somatically mutated driver genes provide a set of gold standard mutations with somatic mutation roles that can be used to assess the performance of OncoMerge. The gold standards are classified by whether the somatic mutation of a gene was cancer-specific or not. The TCGA consensus^6^ and Cancer Gene Census (CGC) from COSMIC^11^ were used to develop gold standards with cancer-specific somatically mutated gene roles. The 20/20 rule^3^, OncodriveROLE^13^, and Tokheim ensemble^12^ were used to create gold standards with somatically mutated gene roles. Comparisons of somatic mutation role between OncoMerge and the gold standards were facilitated by converting oncogenes to Act and tumor suppressors to LoF. Finally, a combined gene role agnostic gold standard was developed based on a union of all somatic mutations from all five gold standards. These gold standards were used to assess the utility of filters and the quality of the OncoMerge integrated somatic mutation matrices through their ability to recall somatic mutations with the appropriate gene role. We chose five gold standards that employed different algorithms for somatic mutation discovery to avoid overfitting to any one gold standard when assessing the performance of OncoMerge.

OncoMerge is designed to construct a comprehensive somatic mutation profile that increases the power to link mutations with cancer phenotypes. Previously, we have used the Systems Genetics Network AnaLysis (SYGNAL) pipeline^19^ to build causal and mechanistic gene regulatory networks (GRNs) for 31 cancers from the TCGA Pan-Cancer Atlas^20^. Using SYGNAL, we linked somatic mutations through transcription factor (TF) and miRNA regulators to the hallmarks of cancer^21, 22^, thereby linking somatic mutations to cancer phenotypes. We hypothesize that integrated somatic mutation matrices from OncoMerge will increase our power to infer causal relationships for pan-cancer SYGNAL networks and that these will yield novel biological insights.

## Results

### Establishing a baseline for the integration of somatic mutations

We developed OncoMerge as a systematic method to integrate PAM, fusion, and CNA somatic mutations into a more comprehensive mutation matrix for subsequent analyses. OncoMerge systematically integrates somatic mutations and defines a role for each gene (**Figure 1**): PAM, fusion, CNA deletion (CNAdel), CNA amplification (CNAamp), Activating (Act), and Loss of Function (LoF). The role assigned to a gene describes the rubric used to integrate the data from the source data matrices.

A significant part of developing OncoMerge was constructing and optimizing the statistical filters that provide an essential quality control step to identify integrated somatically mutated genes more likely to be functional in tumor biology. The selection and optimization of OncoMerge statistical filters were performed using the 9,584 patient tumors from 32 cancers profiled by the TCGA Pan-Cancer Atlas^1, 6, 20^ (cancer type abbreviations can be found in STAR Methods). We used three metrics to assess the value of potential filters: 1) impact on the number of somatically mutated genes (**Figure 2A**); 2) impact on the distribution of the number of genes mapping to genomic loci (**Figure 2B**); and 3) significance of the overlap between somatically mutated genes from OncoMerge with gold standard datasets (including overlap with gene roles and tumor-specific gene roles; **Figure 2C**; **Table S2** & **S3**). These metrics ensure that the integrated somatic mutations are consistent with prior knowledge and that the size of CNA mutations does not overwhelm the integration algorithm.

**Figure 2.**
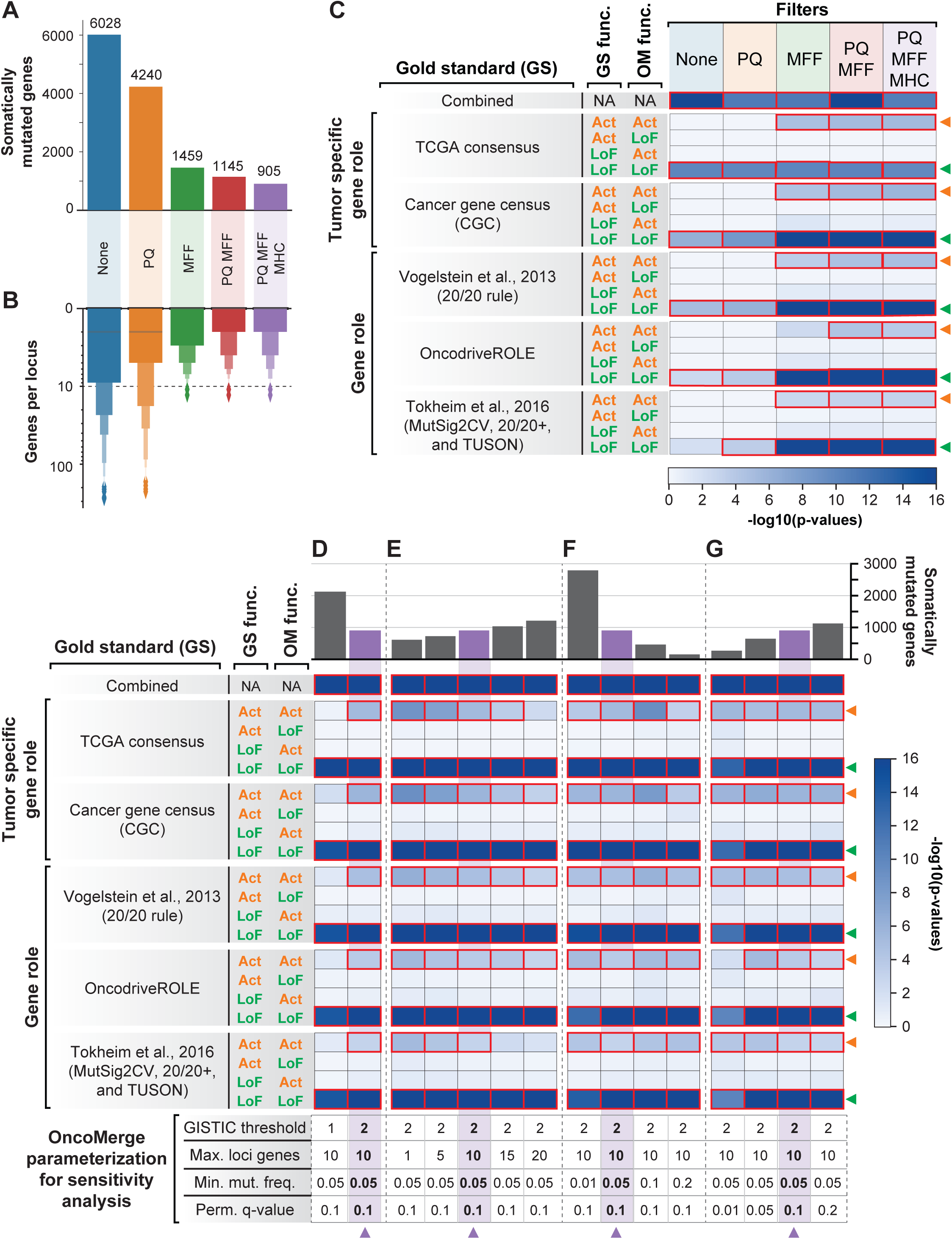
Assessing the performance of OncoMerge filters and conducting sensitivity analyses. **A.** Impact of filter sets on the number of somatically mutated genes inferred by OncoMerge in at least one cancer. **B.** Impact of filter sets on the distribution of genes per CNA locus using the same set of filtering conditions (y-axis is distributed on a log scale). The dashed line indicates the ten genes per loci cutoff that invokes the MFF filter. **C.** Enrichment of the gold standard (GS) Act or LoF somatic mutations with OncoMerge (OM) Act or LoF somatic mutations for each filtering condition: no filters (None); PQ filter; MFF; combined PQ and MFF; and combined PQ, MFF, and MHC. Significant enrichments from **C** are highlighted in red and had p-value less than or equal to the Bonferroni corrected α level of 4.8 x 10^-4^ (α = 0.05, number of tests = 105, Bonferroni corrected α = α / number of tests = 0.05 / 105 = 4.8 x 10^-4^). The orange arrowheads indicate OM Act vs. GS Act, and the green arrowheads indicate OM LoF vs. GS LoF. **D.** Comparing GISTIC thresholds: cutoff of one equates to shallow amplification and deletions, and cutoff of two equates to deep amplifications and deletions. **E.** Comparing the possible values for the MFF filter parameter maximum number of genes in the loci (Max. loci genes) with 1, 5, 10, 15, and 20 genes. **F.** Comparing the possible cutoff values of the minimum mutation frequency (Min. mut. freq.) with 1%, 5%, 10% and 20%. **G.** Comparing the possible cutoff values of the PQ filter permuted q-value (Perm. q-value) with 0.01, 0.05, 0.1 and 0.2. Significant enrichments from **D**-**G** are highlighted in red and had p-value less than or equal to the Bonferroni corrected α level of 2.4 x 10^-3^ (α = 0.05, number of tests per parameters = 21, Bonferroni corrected α = α / number of tests per parameter = 0.05 / 21 = 2.4 x 10^-3^). The purple arrowheads indicate the final parameterization chosen for OncoMerge.

First, we needed to prove that the seed genes alone would not drive significant enrichment for the gold standard comparisons. We applied OncoMerge to somatic mutation matrices with randomized gene labels from the TCGA Pan-Cancer Atlas without any filters applied. None of the gold standards significantly overlapped with the OncoMerge identified somatic mutations (all p-values ≥0.4; **Table S4**). Thus, even though MutSig2CV and GISTIC 2.0 determine the seeds genes, the actual somatic mutation data and filters are required to achieve the full integration potential of OncoMerge. This result from applying OncoMerge to randomized data demonstrates that the signal from the gold standards is not driven by the seed genes alone. Thus we can safely use the gold standards to assess the performance of OncoMerge.

Next, we determined the integration baseline by applying OncoMerge to the TCGA Pan-Cancer Atlas without filtering. Slightly less than one-third of the genome was considered somatically mutated in at least 5% or greater of tumors in at least one of the 32 cancers (30% or 6,028 genes, **Figure 2A**). We observed a significant overlap between OncoMerge somatically mutated genes and the combined gold standard (genes = 395, p-value = 1.1 x 10^-44^, **Figure 2C**) when gene role was not considered. Significant overlaps existed between the LoF somatic mutations from three gold standards (TCGA consensus, CGC, and Vogelstein) with the somatic mutations with the LoF predicted role from OncoMerge (**Figure 2C**). None of the comparisons of Act somatic mutations were significantly overlapping (**Figure 2C**). Many of the 6,028 genes map to the same copy number alteration genomic locus (**Figure 2B**). These unfiltered results reveal two main integration biases. First, the overlaps were insignificant between Act somatic mutations and previously identified Act mutations. Second, the integration with CNAs is causing the inclusion of many passenger mutations mapping to the same genomic locus. OncoMerge applied to the TCGA Pan-Cancer Atlas without filtering provides a baseline to benchmark success. Addressing the integration biases we observed is our impetus for developing and optimizing filters for OncoMerge.

### Developing a filtering strategy for the integration of somatic mutations

The power of integration is that aggregating somatic mutation information can boost the mutation frequency of a gene enough to become significant, even though the constituent mutations do not reach significance alone. The first filter determined if the final mutation frequency after integrating PAM, fusion, and CNA somatic mutations is larger than expected by chance alone. A permutation-based approach empirically determined the background integrated mutation frequency distribution. Then the observed frequencies are compared to the randomized background distribution to calculate permuted p-values, which are corrected using the Benjamini-Hochberg method to provide permuted q-values. A permuted q-value ≤ 0.1 denotes a significant final mutation frequency. The permuted q-value (PQ) filter reduced the number of somatically mutated genes to 4,240 (**Figure 2A**). This filtering improved LoF somatic mutations from three to four gold standards (TCGA consensus, CGC, Vogelstein, and OncodriveROLE) with the somatic mutations that had the LoF predicted role from OncoMerge. Still, the Act comparisons did not show significant enrichment (**Figure 2C**). The PQ filter had a minimal impact on the number of genes per locus (**Figure 2B**). This lack of significant overlap for Act somatic mutations demonstrates that further filtering is required.

A key consideration in developing OncoMerge was that integrating the somatic mutation types should highlight the functional somatic mutations over passenger mutations. Therefore, we created a filter to prioritize somatically mutated genes more likely to be functional. An average CNA encompasses 3.8 ± 7.9 Mb of genomic sequence^23^, and genomic segments of this size typically include many genes. These large genomic regions make it difficult to determine which of the affected genes are the functional gene(s) underlying the CNA locus without integrating additional information. We assert that passenger genes underlying a CNA locus are considered noise and can be identified by the lack of allelic heterogeneity. Thus, functional gene(s) can be identified through allelic heterogeneity that boosts the somatic mutation frequency for a gene above the background CNA frequency. We designed a low-pass filter that retains only the gene(s) with the maximum final frequency (MFF). The MFF filter is only applied if a locus has more than ten genes. Application of the MFF filter dramatically reduced the number of somatically mutated genes from 6,028 to 1,459 (**Figure 2A**) and the number of genes per locus (**Figure 2B**). The MFF filter also helps make the average contribution of PAMs and CNAs to the final mutation frequency more even (**Figure S2**; **Table S5**). We additionally observed a marked improvement in overlap with the gold standards. Significant enrichment was observed for four Act gold standards with somatic mutations that OncoMerge predicts as Act. All five of the LoF gold-standard versus OncoMerge predicted LoF comparisons (**Figure 2C**). Thus, the MFF filter directly addresses the issue of too many genes in a CNA locus. Removing more than three-quarters of the somatically mutated genes improves the overlaps with gold standards.

We then assessed the impact of applying both the PQ and MFF filters. Simultaneous application of both filters reduced the number of somatically mutated genes beyond the MFF filter (1,145 genes; **Figure 2A**), and the number of genes per locus was further improved (**Figure 2B**). There was also an improvement in the significant overlap with gold standards where all five LoF gold-standard versus OncoMerge predicted LoF and significant overlap for four Act gold-standard versus OncoMerge predicted Act (**Figure 2C**). Importantly, none of the gold standard Act versus LoF or LoF versus Act comparisons were significant for any filter combination, demonstrating that the OncoMerge predicted roles are consistent with prior knowledge.

### Reducing biases due to microsatellite instability and hypermutation

Microsatellite instability (MSI) and hypermutation drastically increase the number of somatic mutations in a tumor. The PQ and MFF filters and OncoMerge’s core algorithm rely upon somatic mutation frequency, which is susceptible to confounding by MSI or hypermutation. Fortunately, all TCGA tumors used in this study are characterized for both MSI^5^ and hypermutation^6^ status (**Figure 3A**). We observed a highly significant positive correlation between MSI/hypermutation frequency and the total number of somatic mutations per cancer after integration by OncoMerge (R = 0.68 and p-value = 2.0 x 10^-5^). This strong positive correlation demonstrates that MSI/hypermutation likely inflates the number of somatic mutations discovered by OncoMerge. Therefore, we created the MSI and hypermutation censoring filter (MHC) to exclude these tumors while OncoMerge determines which genes to include in the final somatic mutation matrix. The mutation status for tumors with MSI and hypermutation are included for genes in the final integrated mutation matrix. Applying the MHC filter alongside the PQ and MFF filters reduced the overall number of somatically mutated genes (905 genes; **Figure 2A**) and had minimal impact on the number of genes per locus (**Figure 2B**; **Data S1**). The combined PQ, MFF, and MHC filters decreased the correlation between the MSI/hypermutation frequency (R = 0.48 and p-value = 5.8 x 10^-3^). All ten of the gold standard Act vs. Act and LoF vs. LoF comparisons were significant. These results established that the MHC filter is valuable for removing passenger mutations introduced by tumors with severely increased somatic mutation rates. The PQ, MFF, and MHC filters comprise the default and final OncoMerge filter set. The filters deal with known complications in cancer genetics and ensure that the mutation roles in the integrated matrix are correctly assigned.

**Figure 3.**
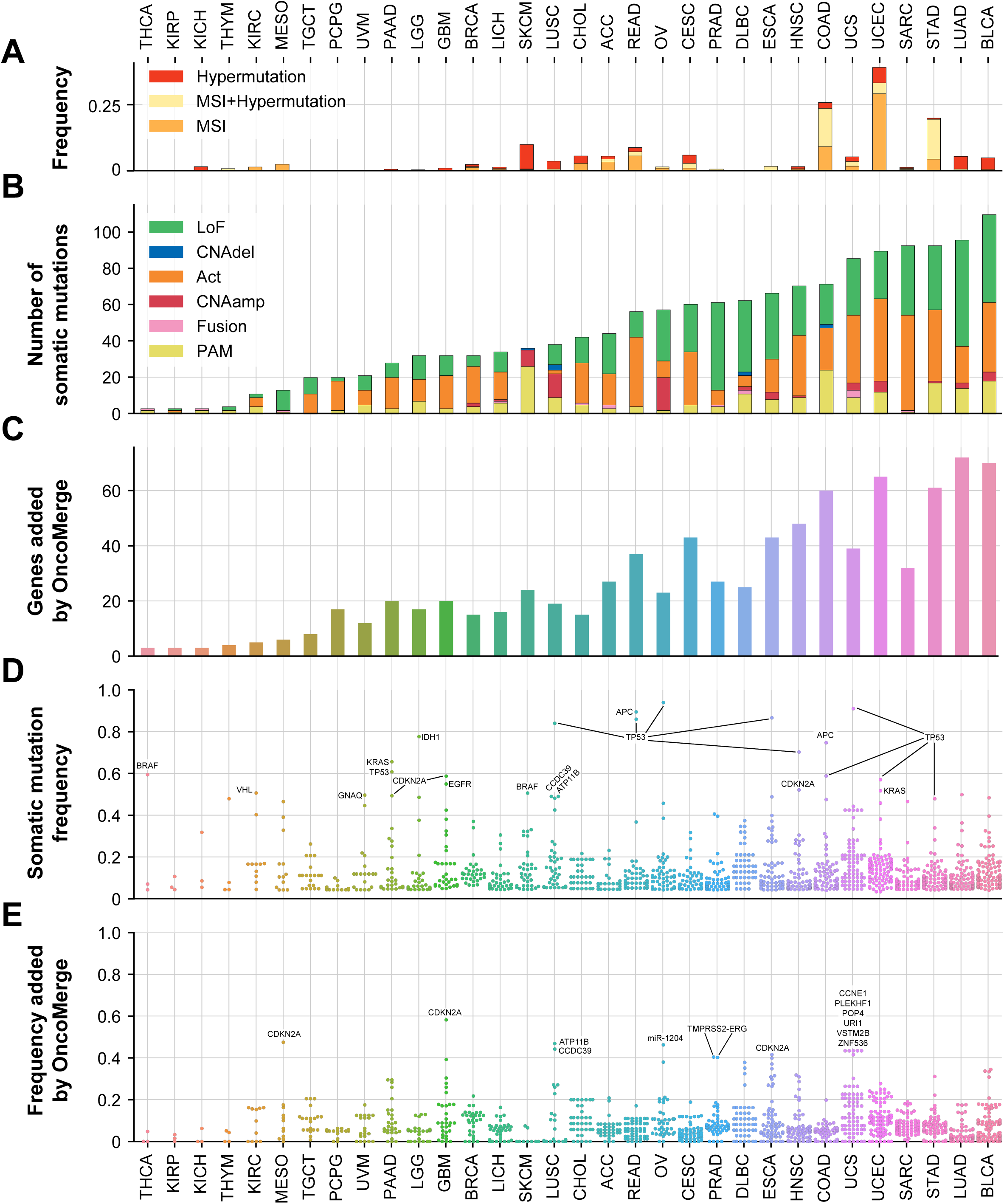
Summary of effect on number and frequency of somatic mutations after integrating mutation types. **A.** Frequency of hypermutation and MSI across cancers. **B**. Number and distribution of mutation types. **C**. Number of somatically mutated genes added because of integration. **D**. Integrated somatic mutation frequencies. **E**. Increases in somatic mutation frequency relative to PAM frequency after integration.

### Sensitivity analyses to optimize filtering cutoffs and parameters

We then used sensitivity analyses of the filtering parameters to determine their optimal parameterization for OncoMerge (**Figure 2D**-**G** and **Table S6**). First, we tested whether a shallow (≥1) or deep (≥2) GISTIC threshold cutoff was optimal for integration purposes (**Figure 2D**). The shallow GISTIC threshold led to many more somatically mutated genes, but impaired the discovery of meaningful activating mutations, as shown by the lack of significant overlap with gold standard activating mutations. Therefore, the deep GISTIC threshold was chosen for OncoMerge. Second, we varied the MFF gene number threshold of a CNA locus across the values 1, 5, 10, 15, and 20 genes (**Figure 2E**). Significant overlap with all gold standard activating mutations is observed up to 10 genes per loci threshold. These results demonstrate that the optimal MFF cutoff is 10 genes per loci. Next, we tested the sensitivity of OncoMerge to the minimum mutation frequency threshold by setting it with the values of 0.01, 0.05, 0.1, and (**Figure 2F**). There was little impact on the significance of the gold-standard analysis across the range of thresholds. However, the number of somatic mutations is significantly impacted by this threshold. A 1% or smaller minimum mutation threshold would be warranted for somatic mutation discovery. On the other hand, ensuring sufficient somatically mutated samples to achieve statistical power for downstream analyses warrants a 5% minimum mutation threshold. Finally, we tested the sensitivity of OncoMerge to the permuted q-value threshold from the PQ filter across the values 0.01, 0.05, 0.1, and 0.2 (**Figure 2G**). The lowest threshold of 0.01 led to a loss of significance for the overlap of the activating mutations for the Vogelstein et al., 2013 gold standard. The number of somatically mutated genes increased by hundreds of genes as the permuted q-value threshold increased, and thus 0.05, 0.1, and 0.2 are all reasonable threshold values. The permuted q-value threshold of 0.1 was chosen because it removed integrated somatic mutations that could have happened by chance alone and retained many somatically mutated genes. These sensitivity analyses provide a reasonable rationale for choosing the values for the filtering cutoffs and parameters, which alternative values might be used, and an idea of which contexts they might be useful.

### Benefits of an integrated somatic mutation matrix

We evaluated the benefits of systematic somatic mutation integration by comparing OncoMerge integrated somatic mutation matrices to those from PAMs. The PAM somatic mutation matrices were used as a reference point because we have successfully used them as the sole source for somatic mutations in previous studies^19, 20^. We assessed the benefits of integration by tabulating the number of somatic mutations and their roles (**Figure 3B**), the number of genes added by integration (**Figure 3C**), and the increase in somatic mutation frequency due to integration (**Figure 3E**). Act and LoF mutations represented the bulk of the somatic mutations in 30 cancers (**Figure 3B**). The THCA and KICH were the only cancers that lacked Act or LoF mutations. Consistent with Agrawal et al. 2014^24^, THCA had only three mutations with a frequency ≥5% BRAF, NRAS, and RET. On the other hand, KICH was under-sampled in the TCGA Pan-Cancer atlas (n = 65), and LoF and Act mutations would likely be discovered with the inclusion of more patient tumors.

We then investigated how many new genes the integration added for each cancer. Integration added at least one somatically mutated gene for each cancer (**Figure 3C**) and more than sixty somatically mutated genes for BLCA, LUAD, STAD, and UCEC (**Figure 3C**). The somatically mutated genes added by OncoMerge make the integrated somatic mutation matrices more comprehensive.

Next, we investigated the frequencies of the somatic mutations from the OncoMerge integrated mutation matrices. The genes with the highest frequency map to well-known oncogenes (e.g., BRAF, KRAS, and EGFR) and tumor suppressors (e.g., APC, CDKN2A, and TP53; **Figure 3D**). The APC gene was mutated in greater than eighty percent of tumors for READ. The TP53 gene was mutated in greater than eighty percent of tumors for ESCA, LUSC, OV, READ, and UCS. These frequently mutated genes in the OncoMerge integrated mutation matrices are consistent with prior knowledge of somatic mutations for each cancer.

Finally, we calculated the frequency added through integration by subtracting the integrated mutation frequency from the PAM frequency. The most substantial increases in somatic mutation frequency were observed for TMPRSS2-ERG in PRAD and CDKN2A in ESCA, GBM, and MESO (**Figure 3E**). Neither TMPRSS2-ERG nor CDKN2A would have been identified as somatically mutated without incorporating fusions and CNAs, respectively. These findings demonstrate that OncoMerge significantly improves the number and frequency of somatically mutated genes in most cancers. Also, these results show that the systematic integration of PAM, fusion, and CNA somatic mutations is crucial for obtaining a comprehensive mutation matrix for each cancer.

### Pan-cancer somatic mutations capture many known tumor suppressors and oncogenes

Genes mutated in multiple cancers are of great interest as selective pressures have found a common solution to influence cancer phenotypes in different contexts. Therefore, we searched for genes somatically mutated in at least five cancers in the OncoMerge integrated mutation matrices. The resulting gene list could be broken down into two groups of somatic mutations: the LoF set (n = 23, **Figure 4A**) and the Act set (n = 13, **Figure 4B**). The genes FBXW7 and KMT2C were somatically mutated with only PAMs. Both genes were previously classified as tumor suppressors^25–27^ and were grouped with the LoF set.

**Figure 4.**
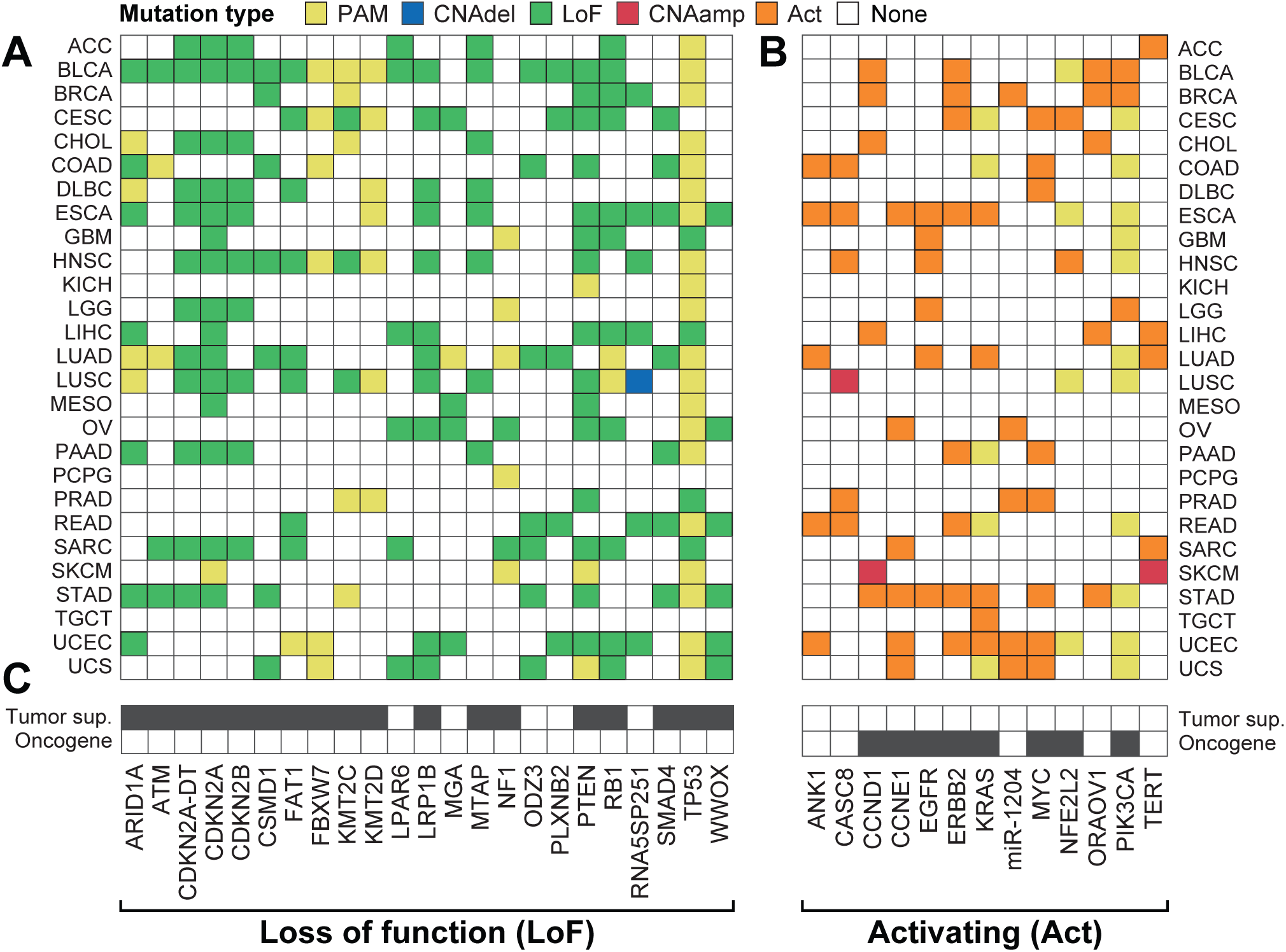
Pan-cancer somatic mutations with a consistent functional impact across at least five cancers. **A**. Pan-cancer somatic mutations from the loss of functions group. **B**. Pan-cancer somatic mutations from the activating group. **C**. Prior knowledge of tumor suppressor or oncogene status for each somatically mutated gene (black square indicates known tumor suppressor or oncogene activity).

The pan-cancer somatically mutated genes harbored many well-known tumor suppressors and oncogenes (**Figure 4C**). As expected, tumor suppressors^27^ were significantly enriched in the LoF group (overlap = 18, p-value = 5.0 x 10^-20^), and oncogenes^28^ were significantly enriched in the Act group (overlap = 8, p-value = 1.6 x 10^-10^). The top three most somatically mutated tumor suppressors were TP53, PTEN, and CDKN2A. These three tumor suppressors control essential checkpoints in the cell cycle, making them functionally interesting. The gene TP53 was somatically mutated in 24 cancers, primarily by PAMs, but four LoFs were also observed for GBM, LIHC, PRAD, and SARC. The top two most mutated oncogenes across cancers were PIK3CA and KRAS, which become overactive kinases when mutated. Both PIK3CA and KRAS have PAM and Act mutation roles across the different cancers, and only the NFE2L2 gene has a similar mixture of PAM and Act mutation roles. The genes CASC8, CCND1, and TERT included cancers with a CNAamp mutation role. The somatic mutation roles are all Act for the remainder of the oncogenes. These pan-cancer analyses further validate the systematic somatic mutation integration by OncoMerge through the unbiased recall of tumor suppressors and oncogenes.

### Improving gene regulatory network inference

Next, the integrated somatic mutation matrices for the TCGA cancer types were used to construct gene regulatory networks (GRNs; **Table 7**) and compared to networks built using only PAMs (legacy)^20^. The GRNs connected somatic mutations to TFs and miRNAs that regulate the expression of a set of genes associated with cancer phenotypes or patient survival.

The average degree was the first metric we considered to compare the GRNs. The degree of a node is the number of edges connecting it to other nodes. The average degree is a standard network metric computed as the average of all node degrees in the network. We found that the average degree was larger for 23 OncoMerge GRNs relative to legacy GRNs (**Figure 5A**). The exceptions were GBM (average degree was equal) and COAD, KIRC, SKCM, and STAD (legacy had a larger average degree). The COAD, SKCM, and STAD cancer types harbor more MSI and hypermutation tumors (**Figure 3A**), and we observed a reduction in the number of COAD and STAD mutations in the OncoMerge GRN relative to the legacy GRN (**Figure 5B**). These results suggest that the MHC filter removed spurious associations. Thus, we have increased the average degree for most networks and addressed a systematic bias in legacy networks.

**Figure 5.**
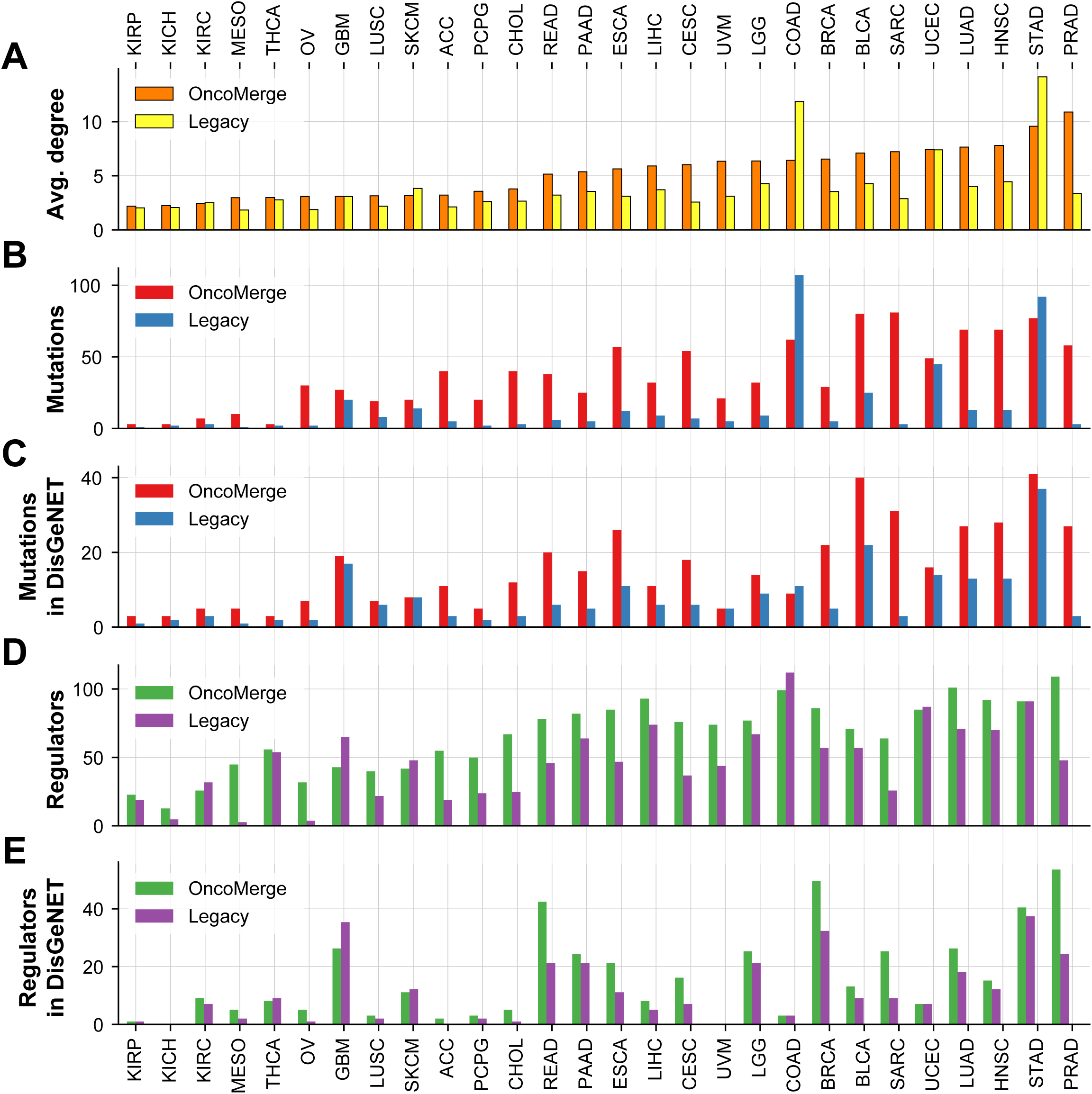
Demonstrating improvements in downstream SYGNAL analysis by comparing GRNs constructed with an OncoMerge integrated somatic mutation matrix versus a legacy network using only PAMs. **A.** Average degree of nodes in the PanCaner SYGNAL networks. **B.** Mutations per cancer network. **C.** Mutations that overlap with genes previously associated with a specific cancer in DisGeNET. **D.** TFs per cancer network. **E.** TFs that overlap with genes previously associated with a specific cancer in DisGeNET.

Next, we compared the number of mutations in each GRN predicted to modulate the activity of regulators. The OncoMerge GRNs contained more somatic mutation nodes than the legacy GRNs for all cancers but COAD and STAD, likely due to MSI and hypermutation (**Figure 5B**). Then, we assessed the recall of somatic mutations previously associated with each cancer from the DisGeNET database^29^. All but two OncoMerge GRNs recalled more previously associated somatic mutations than the legacy GRNs (**Figure 5C**). The exceptions were UVM with the same amount and COAD with fewer (**Figure 5C**). These results demonstrate that OncoMerge integrated mutation matrices provide increased power to infer associations with somatic mutations, especially previously associated with each cancer.

Finally, we considered the number of causal and mechanistic TFs in each GRN. The OncoMerge GRNs contained more predicted TFs than legacy for 22 GRNs, the same number of TF regulators for STAD, and fewer TFs for COAD, GBM, KIRC, THCA, and UCEC (**Figure 5D**). We also assessed the recall of TFs previously associated with each cancer from the DisGeNET database^29, 30^. Twenty of the OncoMerge GRNs recalled more previously associated TFs than legacy GRNs (**Figure 5E**). The COAD and UCEC GRNs had the same amount, and GBM, KIRP, SKCM, and THCA had fewer, and KICH and UVM had no recall of previously associated TFs in either GRN (**Figure 5E**). In summary, using OncoMerge integrated mutation matrices constructs GRNs that are more extensive and biologically meaningful.

### Comparing active and static TF regulatory network architectures

The interactions between TFs are important for generating the transcriptional state of a human cell. The underlying architecture of TF regulatory networks, comprised of TFs and their interactions, are typically explored by enumerating all three-node network motifs and computing their enrichment or depletion into triad significance profiles (TSPs)^31^. Most studies of network motif enrichment have relied upon unsigned interactions^31–36^, which ignore whether the interaction is activating or repressing. To facilitate comparisons, our first analysis of network architecture uses unsigned TSPs to compare static and active TF regulatory networks. Static TF regulatory networks were constructed using chromatin accessibility and DNA binding motifs for 41 cell types^32^. These TF regulatory networks are static because they do not incorporate gene expression data in their construction. Active TF regulatory networks are derived from the OncoMerge augmented SYGNAL pan-cancer GRNs, which were trained using patient tumor transcriptional data and therefore are comprised of active TF regulatory interactions. We calculated TSPs for twenty-five TF regulatory networks and the median TSP (**Figure 6A** & **B**; **Table S8**). We excluded the cancer types DLBC, KICH, KIRP, OV, TGCT, and THYM because they had too few inferred regulatory interactions (< 50 interactions). In addition, we recalculated the TSPs for the static TF regulatory networks using a more recent version of the mfinder algorithm (**Figure 6B**).

**Figure 6.**
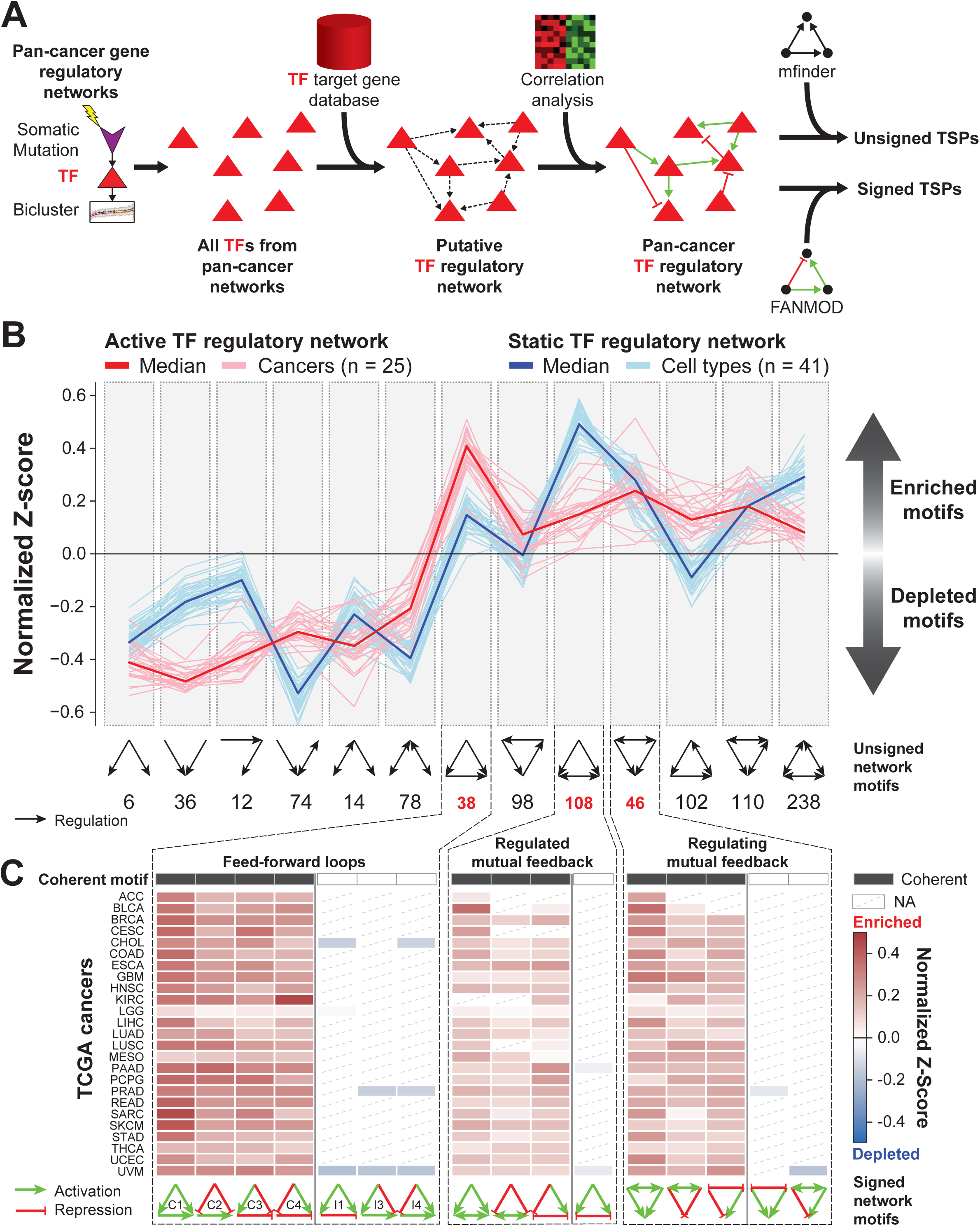
The architecture of functional disease-specific TF regulatory networks from human tumors. **A**. Active TF regulatory network construction pipeline: 1) TFs from all cancer regulatory networks were identified, 2) A putative map of TF regulatory network interactions was constructed, 3) TF → TF relationships were filtered using Pearson’s correlations computed from patient tumor data, and 4) compute the triad significance profiles using mfinder. **B**. Comparison of active TF regulatory network based on SYGNAL GRNs (red) to the static TF regulatory network based on ENCODE DNA binding and accessibility (blue, Neph et al., 2012). **C**. FANMOD enrichment normalized Z-scores for the three most enriched motifs from the active TF regulatory network after incorporating TF regulatory interaction roles (activation or repression). The first row, titled Coherent motifs, is shaded when the motif configuration is coherent and white when it is incoherent. Normalized Z-scores are reported for each cancer, and diagonal dashed lines are inserted when no Z-score was returned. The network motif can be found at the bottom of each column, colored with regulatory roles. C1, C2, C3, C4 = coherent FFLs. I1, I2, I3, I4 = incoherent FFLs.

The median TSPs of the active and static TF regulatory networks were highly correlated (R = 0.75, p-value = 3.0 x 10^-3^; **Figure 6B**). Demonstrating that the architecture of the active network resembles the static network. However, the maximum enriched network motifs were different.

The regulated and regulating feedback motifs (motifs 108 and 46) were the most highly enriched motifs from the static TF regulatory networks and were still enriched, although not as significant, in the active networks. In contrast, the feed-forward loop (FFL, motif 38) is the most highly enriched motif in the active TF regulatory networks. These two motifs are quite similar in structure and differ only by a single edge. Feedback motifs and FFLs can be further broken down into ten and eight signed network motifs that each have a unique functional output^37^. Thus, we can discover what functions are being selected for by evolution in general and the microcosm of tumor biology by exploring the enrichment of signed network motifs.

### Coherent feed-forward loops enriched in active TF regulatory networks

Incorporating the sign of the regulatory interactions (activating or repressing) splits the FFL motif into eight signed network motifs classified as coherent (C1, C2, C3, C4) and incoherent (I1, I2, I3, I4)^37^. Simulation studies have demonstrated that coherent FFLs lead to delays in target gene expression, and incoherent FFLs accelerate target gene expression^37^. FFLs were significantly enriched in active TF regulatory networks, which led us to question whether coherent, incoherent, or both FFLs were enriched. In active GRNs, the sign of the correlation between the TF regulator to TF target can be used to determine the sign of the interaction (R > 0 equates to activation, R < 0 equates to repression). The four coherent FFLs were enriched in the active TF regulatory networks (**Figure 6C**; **Table S9**), and incoherent FFLs were severely under-enriched (Z << 0). In summary, coherent FFLs were enriched in our active TF regulatory networks, suggesting that transcriptional delay mechanisms must provide a valuable function for TF regulatory networks.

### Coherent switch-like feedback motifs enriched in active TF regulatory networks

The regulated and regulating mutual feedback motifs have a two-node feedback loop at their core. The double-positive and double-negative two-node mutual feedback loops act like switches^38^. We tested the twenty signed regulated and regulating mutual feedback network motif configurations for enrichment in TF regulatory networks. Three regulating and three regulated signed mutual feedback motifs (**Figure 6C**; **Table S9**). These six enriched regulated and regulating mutual feedback motifs had a common configuration. Firstly, all the network motifs were coherent. Coherent regulated and regulating feedback loops have interaction signs between the feedback loop that are either double-positive or double-negative. And the regulated or regulating node interacts with the feedback loop nodes using the same sign for double-positive feedback loops and the opposite sign for double-negative feedback loops. Thus, there are three coherent configurations for both regulated and regulating mutual feedback motifs making six total, coinciding with the six enriched configurations (**Figure 6C**; **Table S9**). The enriched motifs containing a double-positive feedback loop had the same interactions with the non-feedback loop node, both activating or repressing (**Figure 6C**). The enriched motif containing a double-negative feedback loop had opposing interactions with the non-feedback loop node, one activating and one repressing (**Figure 6C**). These enriched signed network motifs are the configurations that function as molecular switches^39^. Again, evolution has selected for coherent network motif configurations likely because of their function.

## Discussion

We avoided overfitting while developing and optimizing parameters for OncoMerge in three ways. First, we used five gold standards that use different methods for somatic mutation discovery to avoid overfitting to one specific gold standard. This diversification approach was successful because we observed variable enrichment scores across the gold standards. Secondly, the sensitivity analyses we conducted over a plausible set of parameter values demonstrated the robustness of OncoMerge to different parameterizations. This is important because it shows that OncoMerge has not been parameterized into an anomalous overfit state. Instead, the parameters were chosen based on carefully considered statistical choices and trends in the data. Thirdly, we avoided overfitting to a specific cancer somatic mutation profile by applying and assessing the performance of OncoMerge across thirty-two cancer types. The ability of OncoMerge to be applied to a Pan-Cancer cohort with many different mutation profiles strongly suggests that OncoMerge should be generalizable to new cancer cohorts. We employed all three of these approaches to avoid overfitting and to ensure that OncoMerge could be applied to new datasets without having to tune parameters.

Additionally, we provide sensitivity analyses that can guide users who want to change OncoMerge parameters by observing how specific parameter values impact its performance. For example, the minimum mutation frequency can be set to zero to conduct somatic mutation discovery, providing a more comprehensive list of somatic mutations and their types. In this study, we chose a five percent cutoff for the minimum mutation frequency to ensure there were enough somatically mutated tumors to power downstream GRN inference. The sensitivity analyses of can be used to guide the choice of OncoMerge parameters to achieve different goals than the default parameterization.

The construction of active GRNs enabled the exploration of signed network motifs and led to the discovery that specific signed network motif configurations are being enriched. The SYGNAL GRN construction method identifies active gene regulatory interactions by discovering interactions that are supported by gene expression data from patient tumors^19^. On the other hand, prior networks were static maps of DNA binding sites constructed using digital genomic footprinting and the similarity of the underlying sequence of the footprints for known DNA binding motifs^32^. The active networks use a correlation-based method to determine TF regulatory roles (activator or repressor) for the interactions, which is not possible using static binding maps. Analyzing signed network motifs provides a leap forward in understanding how the underlying architecture of GRNs functions in real-world biological systems. OncoMerge integrated somatic mutations offer a more solid platform to infer active GRNs that can be used to explore the functional architecture of TF regulatory networks.

We discovered that coherent regulated and regulating feedback and FFL network motifs were enriched in cancer TF regulatory networks. We cannot say whether this enrichment of network motifs will generalize to all active GRNs or if this is a cancer-specific phenomenon. In normal organismal development, feedback motifs have been previously shown to be essential for cell fate decision-making^40, 41^. On the other hand, in tumor cells and other cells in the tumor microenvironment, the enriched feedback motifs may be maintaining a cell fate, or the disease could be coopting the circuit to drive tumor biology. Likewise, coherent FFL network motifs have also been associated with enhanced drug resistance^42^. These coherent motifs are relevant for normal and diseased cell biology, and evolution has specifically selected these motif configurations because of their unique functional outputs.

Future improvements to the OncoMerge algorithm include a more quantitative integration approach for the somatic mutations, a replacement for or an improved maximum final frequency filter, aggregation across pathways, and a determination of whether other genomic features may be integrated (ecDNA^43^ or epigenomics^44^). Additionally, in future single-cell studies with both transcriptome and genome information, it would be helpful to have an OncoMerge implementation that integrates PAM, fusion, and CNA for every single cell. We envision OncoMerge as a valuable tool in the somatic mutation characterization pipeline. We hope it will facilitate multi-omic studies and lead to novel discoveries that can be translated into clinical insights.

### Limitations of the study

Currently, OncoMerge assumes that the somatic mutations will be either PAM, CNA, or gene fusions, meaning it will miss somatic mutations such as ecDNA^43^, epigenomics^44^, etc. Somatic mutations were seeded by PAMs or CNAs that were mutated more than expected by chance alone, which may exclude mutations of lower frequency from being discovered. Future studies could be used to come up with alternative methods of seeding somatic mutations. Using a five percent cutoff for somatic mutation frequency means that lower-frequency mutations will be overlooked. Setting the minimum mutation frequency cutoff to less than five percent would provide a complete list of somatic mutations.

## STAR Methods

### Resource availability

#### Lead contact

Requests for further information should be directed to the lead contact, Christopher Plaisier.

#### Materials availability

This study did not generate new materials.

#### Data and code availability

This paper analyzes existing, publicly available data. All the datasets used as input for our study were deposited in Figshare (https://doi.org/10.6084/m9.figshare.21760964.v1) and they are publicly available as of the date of publication. New datasets generated from our studies were deposited in Figshare (https://doi.org/10.6084/m9.figshare.20238867.v1) and they are publicly available as of the data of publication.

The OncoMerge original code has been deposited at GitHub (https://github.com/plaisier-lab/OncoMerge) and is also accessible through Zenodo using the DOI https://doi.org/10.5281/zenodo.5519663.

Any additional information required to reanalyze the data reported in this paper is available from the lead contact upon request.

##### TCGA cancer abbreviations

Study Abbreviation: Study Name
ACC: Adrenocortical carcinoma
BLCA: Bladder Urothelial Carcinoma
LGG: Brain Lower Grade Glioma
BRCA: Breast invasive carcinoma
CESC: Cervical squamous cell carcinoma and endocervical adenocarcinoma
CHOL: Cholangiocarcinoma
COAD: Colon adenocarcinoma
ESCA: Esophageal carcinoma
GBM: Glioblastoma multiforme
HNSC: Head and Neck squamous cell carcinoma
KICH: Kidney Chromophobe
KIRC: Kidney renal clear cell carcinoma
KIRP: Kidney renal papillary cell carcinoma
LIHC: Liver hepatocellular carcinoma
LUAD: Lung adenocarcinoma
LUSC: Lung squamous cell carcinoma
DLBC: Lymphoid Neoplasm Diffuse Large B-cell Lymphoma
MESO: Mesothelioma
OV: Ovarian serous cystadenocarcinoma
PAAD: Pancreatic adenocarcinoma
PCPG: Pheochromocytoma and Paraganglioma
PRAD: Prostate adenocarcinoma
READ: Rectum adenocarcinoma
SARC: Sarcoma
SKCM: Skin Cutaneous Melanoma
STAD: Stomach adenocarcinoma
TGCT: Testicular Germ Cell Tumors
THYM: Thymoma
THCA: Thyroid carcinoma
UCS: Uterine Carcinosarcoma
UCEC: Uterine Corpus Endometrial Carcinoma
UVM: Uveal Melanoma

## Method details

### Clinical and molecular data from TCGA

These studies used standardized, normalized, batch corrected, and platform-corrected multi-omics data generated by the Pan-Cancer Atlas consortium for 11,080 participant tumors^20^. Complete multi-omic profiles were available for 9,584 patient tumors. TCGA aliquot barcodes flagged as “do not use” or excluded by pathology review from the Pan-Cancer Atlas Consortium were removed from the study. The overall survival (OS, OS.time) data used were obtained from Liu et al. 2018^45^.

- <u>Somatic protein affecting mutations (PAMs) in TCGA</u> – Somatic PAMs were identified by the Multi-Center Mutation Calling in Multiple Cancer (MC3) project^1^ and were downloaded from the ISB Cancer Gateway in the Cloud (ISB-CGC; https://isb-cgc.appspot.com/). PAMs were required to have a FILTER value of either: PASS, wga, or native_wga_mix. In addition, all PAMs needed to be protein-coding by requiring that Variant_Classification had one of the following values: Frame_Shift_Del, Frame_Shift_Ins, In_Frame_Del, In_Frame_Ins, Missense_Mutation, Nonsense_Mutation, Nonstop_Mutation, Splice_Site, or Translation_Start_Site. Additionally, mutation calls were required to be made by two or more mutation callers (NCALLERS > 1). When both normal tissue and blood were available, the blood was used as the germline reference.
- <u>Statistical significance of PAMs in TCGA</u> – The likelihood that a gene is somatically mutated by chance alone was determined using MutSig2CV^7^ and downloaded for each cancer from the Broad GDAC FIREHOSE (https://gdac.broadinstitute.org/). Genes with a MutSig2CV False Discovery Rate (FDR) corrected p-value (q-value) less than or equal to 0.1 were considered significantly mutated^7^.
- <u>Somatic transcript fusions in TCGA</u> – The TumorFusions portal^2^ provides a pan-cancer analysis of tumor transcript fusions in the TCGA using the PRADA algorithm^8^.
- <u>Somatic copy number alterations (CNAs) in TCGA</u> – Genomic regions that were significantly amplified or deleted were identified using Genomic Identification of Significant Targets in Cancer (GISTIC2.0)^9^ and downloaded for each cancer from the Broad GDAC FIREHOSE.

### Somatic mutation data import and preprocessing

An essential first step in OncoMerge is loading up and binarizing the somatic mutation data (**Figure S1**). The somatic mutation data comprised of four primary matrices: 1) PAMs, 2) fusions, 3) CNA amplifications (CNAamps), and 4) CNA deletions (CNAdels) (**Figure 1**). In addition, two derivative matrices Act and LoF are created by merging the PAM with the CNAamps or CNAdels matrices, respectively (**Figure 1**). All files are formatted as comma-separated values (CSV) files with genes as rows and patients as columns unless otherwise noted.

- <u>PAM matrix</u> - The matrix values are [0 or 1]: zero indicates the gene is not mutated in a patient tumor, and one indicates the gene is mutated in a patient tumor.
- <u>Fusion matrix</u> - The matrix values are [0 or 1]: zero indicates no gene fusion in a patient tumor, and one indicates the gene fused to another genomic locus in a patient tumor.
- <u>CNAamp and CNAdel matrices</u> – The all_thresholded_by_genes.csv GISTIC output file is used to populate the CNAamp and CNAdel matrices. The all_thresholeded_by_genes matrix values range from -2 and have no positive bound, and the values indicate the copy number relative to the background. A cutoff of greater than or equal to 2 was used to identify deep amplifications and less than or equal to -2 for deep deletions. Only deep amplifications or deletions were included in these studies due to heterogeneity of cell types and tumor biopsy purity. Oncomerge allows this threshold to be modified through a command line parameter (’-gt’ or ’--gistic-threshold’).
  - <u>CNAamp matrix</u> – The matrix values are [0 or 1]: zero indicates a gene is not amplified in a patient tumor, and one indicates the gene is amplified in a patient tumor.
  - <u>CNAdel matrix</u> – The matrix values are [0 or 1]: zero indicates a gene is not deleted in a patient tumor, and one indicates a gene is deleted in a patient tumor.
- <u>Act matrix</u> – The Act matrix is the bitwise OR combination of the PAM, Fusion, and CNAamp matrices. The Act matrix has genes as rows and patients as columns. The matrix values are [0 or 1]: zero indicates the gene is not mutated or amplified in a patient tumor, and one indicates the gene is either mutated, fused, amplified, or some combination in a patient tumor.
- <u>LoF matrix</u> – The LoF matrix is the bitwise OR combination of the PAM, Fusion, and CNAdel matrices. The LoF matrix has genes as rows and patients as columns. The matrix values are [0 or 1]: zero indicates the gene is not mutated or deleted in a patient tumor, and one indicates the gene is either mutated, fused, deleted, or some combination in a patient tumor.

### Seeding OncoMerge with putative somatic mutations

OncoMerge focuses on likely causal somatic mutations by considering only somatic mutations that were statistically shown to be mutated more often than expected by chance alone. Likely causal somatic mutations are also required to have a mutation frequency greater than 5%, the definition of a common mutation^46^, as this ensures sufficient patient tumors will be mutated to power downstream analyses. These statistically significant common mutations were used as seeds for OncoMerge integration. PAMs used as seeds were identified with MutSig2CV q-values less than or equal to 0.1^14^ and a mutation frequency greater than 5%. Gene fusions used as seeds were identified as significant in PRADA^2, 8^ and had a mutation frequency greater than 5%. CNAamps or CNAdels used as seeds were identified as significantly amplified or deleted from the amplified genes (amp_genes) or deleted genes (del_genes) GISTIC output files with residual q-values less than or equal to 0.05^47^. CNAs from sex chromosomes (X and Y) were excluded. Genes from sex chromosomes can enter OncoMerge as seeds from PAMs or fusions. These seed genes become the starting point of the OncoMerge integration. Subsequent steps determine if Act or LoF merged mutation profiles or their component PAM, Fusion, CNAamp, or CNAdel mutation roles are the most appropriate integration model for a gene.

### Merging somatic mutations in OncoMerge

The mutation role for each seed gene is assigned based on the frequencies of the mutation types for a gene from the original (PAM, Fusion, CNAamp, CNADel) and merged (Act and LoF) somatic mutation matrices and statistical thresholds for PAM (MutSig2CV) and CNAs (GISTIC). The function c) is applied to each seed gene to choose the mutation role using the following parameters: *f_MMF_* the minimum mutation frequency (defaults to 5%), *f_Act_* the frequency of the merged Act mutations, *f_LoF_* the frequency of the merged LoF mutations, *f_PAM_* the frequency of the PAM mutations, *f_Fusion_* the frequency of the gene fusion mutations, *f_CNAamp_* the frequency of CNA amplification mutation, *f_CNAdel_* the frequency of the CNA deletions mutations, *qv_MutSig2CV_* significance of PAM mutations as MutSig2CV q-value, and *qv_GISTIC_* significance of CNA mutations as GISTIC residual q-value.

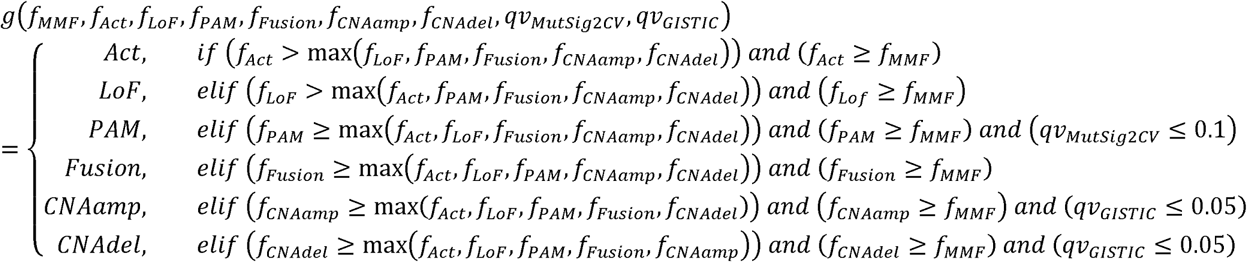

The mutation role for each seed gene is chosen using this decision tree based on mutational frequencies and statistical significance. The Act and LoF are first in the decision tree because merging mutation types should lead to a larger mutation frequency than any individual source mutation frequency (*f_LoF_, f_PAM_, f_Fusion_, f_CNAamp_, f_CNAdel_*). A strict inequality (greater than) is used so mutation frequency as a source mutation frequency. If an Act or LoF integrated mutation role is not chosen, then the source mutation with the highest frequency is chosen. And in the case of ties the non-strict inequalities (greater than or equal to) determine the order of preference for the tied mutational roles: PAM > Fusion > CNAamp > CNAdel. This ordering ensures that integrated mutation roles are chosen when possible and that the most frequent source mutation role is otherwise chosen. The PQ, MFF, and MHC filters further modify the assigned gene mutation roles to determine the final gene mutation role.

### Permuted q-value (PQ) filter

For putative Act and LoF mutations, a permuted q-value is computed by randomizing the order of rows in the PAM, Fusion, and CNA mutation matrices’ and then calculating the randomized frequency distribution for Acts and LoFs. The observed frequency for an Act or Lof mutation is then compared to the randomized frequency distribution to compute the permuted p-value. Permuted p-values are corrected into q-values using the multiple-test Benjamini-Hochberg FDR-based correction method. Only Acts or LoFs that had a permuted q-value ≤ 0.1 were retained. Any Act or LoF with a permuted q-value > 0.1 was set to the mutation role of either PAM, Fusion, CNAamp, or CNAdel based on which mutation role had the highest frequency. This modifies the function *g* into *g_PQ_* that includes the permuted q-value as a new input variable *qv_permuted_*, and is included as a constraint for the calls of Act and LoF.

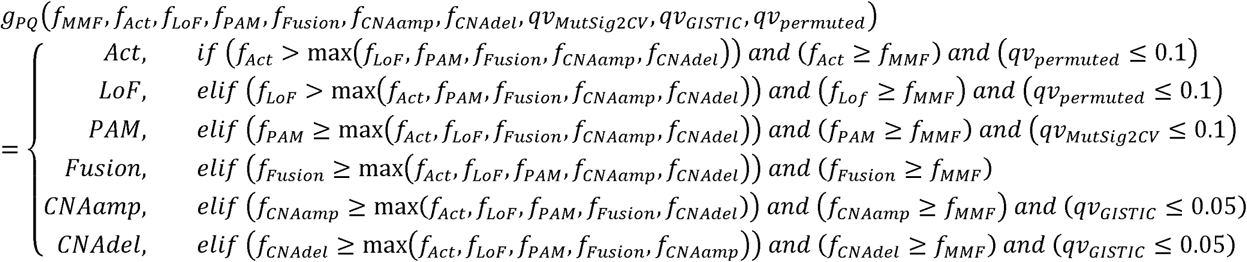

The permuted q-value cutoff defaults to 0.1 and can be set to another value through a command line parameter (’-pq’, --perm_qv’).

### Maximum final frequency (MFF) filter

The maximum final frequency (MFF) filter is a low-pass genomic filter designed to remove passenger genes from frequently mutated CNA loci that contain many underlying genes. By default, the filter is applied when there are 10 or more genes in a locus. Let *L_MFF_* be the set of loci with greater than 10 genes, *locus* be defined as the set of genes in a CNA locus, and *len* a function that returns the number of genes in a set.

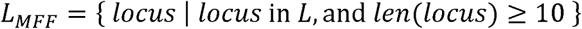

Let *G_locus_* be the set of all genes underlying a CNA locus.

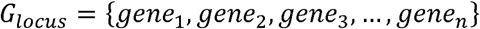

The first step of the filter defines the maximum mutation frequency (*f_MFF_*) for the genes of *locus*. This requires using two functions: *f_req_* which returns the mutation frequency for a gene, and *max* which returns the maximum value from a set.

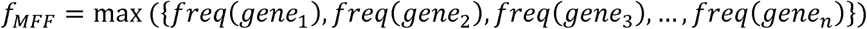

Only the gene(s) that have a mutation frequency equal to the *f_MFF_* are retained for *locus_MFF_*.

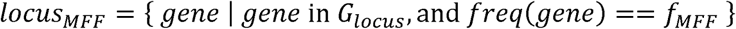

The genes from each *locus_MFF_* are included in the final mutation matrix. The number of genes underlying a CNA locus can be set through a command line parameter (’-mlg’, -- max_loci_genes’).

### Microsatellite hypermutation censoring (MHC) filter

The TCGA tumors used in this study have been characterized for both MSI^5^ and hypermutation^6^ (**Table S1**). The tumors with MSI or hypermutation are loaded as a blocklist of patient IDs through a command line parameter (’-bl’ or ’--blocklist’). All tumors in the blocklist are excluded from consideration by the PQ and MFF filters while determining the genes to include in the final somatic mutation matrix. The mutation status for blocklist tumors are included in the final integrated mutation matrix.

### Optimizing OncoMerge filtering parameters

Sensitivity analyses of filtering parameters (GISTIC threshold, maximum loci genes, minimum mutation frequency, and permuted q-value cutoff) for the OncoMerge algorithm was conducted by varying one input parameter while fixing all others. The number of somatically mutated genes and enrichment of gold standards from each parameterization of OncoMerge was evaluated to determine the optimal values for each input parameter.

### OncoMerge outputs

OncoMerge provides four output files that provide valuable information about the integration process and the final integrated mutation matrix that can be used in downstream studies. Here is a brief description of each file and its contents:

- <u>oncoMerge_mergedMuts.csv</u> – The integrated mutation matrix is comprised of genes (rows) by patient tumors (columns) of mutation status after integration by OncoMerge. The matrix values are [0 or 1]: zero indicates that the gene is not mutated in a patient tumor, and one indicates that the gene was mutated in a patient tumor.
- <u>oncoMerge_CNA_loci.csv</u> – A list of the genes mapping to each CNAamp or CNAdel locus included in the OncoMerge integrated mutation matrix.
- <u>oncoMerge_ActLofPermPV.csv</u> – List of all significant Act and LoF genes, their OncoMerge mutation role, frequency, empirical p-value, and empirical q-value. This output is before the application of the low-pass frequency filter.
- <u>oncoMerge_summaryMatrix.csv</u> – Matrix of genes (rows) by all information gathered by OncoMerge.

To aid in comparisons between runs, we provide the save permutation option (’-sp’ or ’-- save_permutation’) to output permutation results so that the same permuted distribution can be used with different parameters in separate runs. We also provide the load permutation option (’-lp’ or ’--load_permutation’) to load up the permuted distribution from a previous run. The permuted distributions are saved in the following files if requested:

- <u>oncomerge_ampPerm.npy, oncomerge_delPerm.npy</u> – Snapshot of the non-deterministic permutation results from combining PAM, Fusion, and CNAamp or PAM, Fusion, and CNAdel frequencies, respectively.

### Gold standard cancer-specific gene role validation datasets

Gold standard datasets are vital to validating the usefulness of each feature in OncoMerge. Two different sources of gold standard cancer-specific gene role (Act or LoF) datasets were used to validate the OncoMerge predicted tumor-specific gene roles:

- <u>TCGA consensus</u>: The TCGA consensus is a list of driver genes identified from the TCGA Pan-Cancer Atlas labeled with somatic mutation role (oncogene or tumor suppressor) and cancer type. The TCGA consensus was constructed by Bailey et al., 2018 wherein they catalog a list of 299 unique oncogenesis associated genes^6^. In the TCGA consensus 280 cancer-specific oncogene roles were identified, and 417 cancer-specific tumor suppressor roles were identified (**Table S2**).
- <u>Cancer Gene Census (CGC)</u>: The CGC from COSMIC is an expert-curated database of human cancer driver genes labeled with somatic mutation role (oncogene and tumor suppressor) and cancer type. The CGC was developed by Catalogue of Somatic Mutations in Cancer (COSMIC) as an expert-curated database of human cancer-driving genes^11^. CGC cancers were mapped to the TCGA cancers by manual curation (**Table S2**). In the CGC 205 cancer-specific oncogene roles were identified, and 304 cancer-specific tumor suppressor roles were identified (**Table S2**).

### Gold standard gene role validation datasets

Three different sources of gold standard gene role (Act or LoF) datasets were used to validate the OncoMerge predicted gene roles:

- <u>20/20 rule</u>: The 20/20 rule defines oncogenes (Act) by requiring >20% of mutations in recurrent positions, and tumor suppressors (LoF) as >20% of recorded mutations are inactivating (missense or truncating)^3^. With the 20/20 rule, 54 oncogene roles were identified, and 71 tumor suppressor roles were identified (**Table S2**).
- <u>OncodriveROLE</u>: OncodriveROLE is a machine learning algorithm that classifies genes according to their role (Act or LoF) based on well-curated genomic features^13^. With OncodriveROLE, 76 oncogene (Act) roles were identified, and 109 tumor suppressor (LoF) roles were identified (**Table S2**).
- <u>Tokheim Ensemble</u>: Ensemble-based method from Tokheim et al., 2016^12^, which integrates MutSigCV, 20/20+, and TUSON methods for predicting gene roles (oncogene and tumor suppressor). With the Tokheim Ensemble, 78 oncogene (Act) roles were identified, and 212 tumor suppressor (LoF) roles were identified (**Table S2**).

### Computing overlap between OncoMerge and gold standards

A hypergeometric enrichment statistic was used to compute the significance of overlap observed between each gene role in OncoMerge versus the gold standards. When possible, the tumor specificity of the gene role was taken into consideration (TCGA consensus and CGC). A total of 105 hypergeometric enrichment tests were conducted for the comparison to gold standards to test out different filters (5 combined gold standard tests + 5 gold standard datasets * 5 filter conditions [None, PQ, MFF, PQ MFF, and PQ MFF MHC] * 4 GS & OM functional tests [Act vs. Act, Act vs. LoF, LoF vs. Act, and LoF vs. LoF] = 105 tests). An α level of 0.05 was chosen, and significant overlaps were determined as p-values less than or equal to the Bonferroni multiple hypothesis corrected alpha level of 4.8 x 10^-4^ (α / number of tests = 0.05 / 105 = 4.8 x 10^-4^). This cutoff ensures that the comparisons to the gold standards are not likely to have occurred by chance alone, even though we conducted 105 independent tests.

For each sensitivity analysis we conducted 21 tests against the gold standards (1 combined gold standard test + 5 gold standard datasets * 4 GS & OM functional tests [Act vs Act, Act vs LoF, LoF vs Act, and LoF vs LoF] = 21 tests). An α level of 0.05 was chosen, and significant overlaps for sensitivity analyses across the potential parameter values for OncoMerge were determined as p-values less than or equal to the Bonferroni multiple hypothesis corrected alpha level of 2.4 x 10^-3^ (α / number of tests = 0.05 / 21 = 2.4 x 10^-3^). This cutoff addresses the impact of the 21 independent tests for each parameter value in the sensitivity analysis.

### Availability of OncoMerge

We provide the Oncomerge software and data in several standard distribution formats to facilitate future studies that aim to integrate somatic mutations. The source code for OncoMerge is available on GitHub (https://github.com/plaisier-lab/OncoMerge). Finally, an OncoMerge Docker image was created that can be run as a virtual machine with all dependencies pre-installed (https://hub.docker.com/r/cplaisier/oncomerge). Detailed documentation is provided, along with a tutorial that describes the use of OncoMerge. The goal of disseminating OncoMerge in these ways is to give end-users flexibility to choose what distribution method best fits their computational platform.

### OncoMerge TCGA Pan-Cancer Atlas input and output files

We also provide the Pan-Cancer Atlas TCGA somatic mutation data used as input for OncoMerge (https://doi.org/10.6084/m9.figshare.21760964.v1). And the resulting OncoMerge integrated somatic mutation matrices for those planning studies that use somatic mutations from the TCGA Pan-Cancer Atlas (https://doi.org/10.6084/m9.figshare.20238867). These integrated somatic mutation matrices can be used for any downstream analyses incorporating somatic mutations and will provide the same power boost observed in our studies. In addition, we also offer the pan-cancer SYGNAL GRNs and TF regulatory networks as supplementary tables (**Tables S7**–**S9**) to expedite systems genetics studies of TCGA cancers.

### TCGA Pan-Cancer SYstems Genetics Network AnaLysis (SYGNAL)

The mRNA and miRNA expression data required to run SYGNAL were obtained from Thorsson et al., 2018^20^. The SYGNAL pipeline is composed of 4 steps and command-line parameters for all programs are described in detail in Plaisier et al., 2016^19^. Each cancer was run separately through the pipeline to reduce the confounding from tissue of origin differences. Highly expressed genes were discovered for each cancer by requiring that genes have greater than or equal to the median expression of all genes across all conditions in ≥ 50% of patients^19^. These gene sets were then used as input to SYGNAL to construct the gene regulatory networks (GRNs) for each cancer.

The underlying cMonkey2 biclustering results are identical to those from Thorsson et al., 2018^20^ as they do not rely upon genetic information. All immune-specific filters were removed for these analyses, and all bicluster filtering was done as described in Plaisier et al., 2016^19^. Using Network Edge Orienting (NEO)^48^ somatic mutations are integrated with bicluster and regulator expression in the next step. Two networks were constructed by applying systems genetics analysis with NEO to the biclusters: 1) GRNs were inferred using PAM-only somatic mutation matrices as a baseline; 2) GRNs were inferred using OncoMerge integrated somatic mutation matrices. Importantly, the PAM-only somatic mutation matrices used were the same ones used as input for OncoMerge.

### TF regulatory network construction for PanCan-SYGNAL networks

A TF regulatory network was built for each cancer in three steps (**Figure 6A**). First, the TFs regulating survival-associated biclusters were extracted from each cancer’s SYGNAL GRN. Second, a preliminary TF_regulator_ TF_target_ regulatory network was constructed based on the presence of a binding site for a putative TF_regulator_ in the promoter of a TF_target_ from the Transcription Factor Target Gene Database^19^ (http://tfbsdb.systemsbiology.net). TF family expansion^19^ was used to supplement TFs that did not have an experimentally determined DNA recognition motif in the database. The assumption was that the motifs within a TF family would not vary significantly. Therefore, TF family members from the TFClass database^49^ with a known DNA recognition motif can be used as a proxy for a TF with no known DNA recognition motif.

Finally, the putative TF_regulator_ →TF_target_ regulatory network was filtered by requiring a significant Pearson correlation between the mRNA expression of the TF_regulator_ and TF_target_ (Pearson’s |R| ≥0.3 and p-value ≤0.05; **Figure 6A**; **Table S9**). The sign of the correlation coefficient can be used to determine the role of a regulatory interaction: a positive correlation coefficient equates to the TF_regulator_ being an activator, and a negative correlation coefficient equates to the TF_regulator_ being a repressor. Networks with fewer than 50 interactions were not included in the analyses as they were not sufficiently powered to run the network motif analysis. The cancer regulatory networks for DLBC, KICH, KIRP, OV, TGCT, and THYM were excluded from further studies.

### TF regulatory network motif analysis

Three-node network motifs were enumerated from the TF regulatory networks using mfinder^50^ in the same manner as Neph et al., 2012^32^ and used to compute triad significance profiles (TSPs)^31^. The parameters used with mfinder v1.20 were^32^: motif size set at 3 (-s 3), requested 250 random networks to be generated (-r 250), and the Z-score threshold was set at -2000 to ensure all motifs are reported (-z -2000). All Z-scores were extracted for each cancer and converted to triad significance profiles using the methods of Milo et al., 2004^31^.

For consistency, the TF regulatory networks for the 41 different cell types from Neph et al., 2012 were downloaded from http://www.regulatorynetworks.org/ and analyzed using the same approach described above.

### Signed network motif analysis incorporating TF regulator interaction roles

The enrichment of signed feed-forward loops (FFLs), regulated feedback, and regulating feedback network motifs was computed using FANMOD^51^, which takes into consideration TF regulatory roles (activation and repression). The command line version of FANMOD from IndeCut^52^ was used with default parameters, except for the inclusion of regulatory role (colored edges)^51^ (fanmod 3 100000 1 <INPUT_FILE> 1 0 1 2 0 1 0 1000 3 3 <OUTPUT_FILE> 1 1). Z-scores for signed FFLs, regulated feedback, and regulating feedback network motifs were extracted for each cancer and converted to triad significance profiles using the methods of Milo et al., 2004^31^. The signed FFL network motifs are broken down into C1, C2, C3, C4, I1, I2, I3, and I4, as described previously^37^.

## Quantification and statistical analysis

A nominal alpha value (p-value or q-vaule cutoff) of 0.05 was used unless otherwise stated. Statistical analyses are described in detail in the methods sections where they were used, and we provide a brief synopsis of the statistical methods below. Hypergeometric enrichment analysis was used to identify significant overlaps of OncoMerge-derived gene sets with gold-standards gene sets. When appropriate for the gold-standard analyses, Benjamini-Hochberg FDR multiple hypothesis correction was applied to the hypergeometric p-values. Cutoff were as described in the methods or results. Permuted p-values were computed for each integrated somatic mutation and Benjamini-Hochberg FDR multiple hypothesis correction was applied to generate permuted q-values. Pearson correlations were used to compare two sets of quantitative values (e.g., number of somatic mutations and MSI/hypermutation frequency) and the correlation coefficient (R) and p-value are reported. Triad significance profiles (TSPs) were used to quantify the enrichment of three node network motifs.

## Supplementary tables

**Table S1.** Blocklist of MSI and hypermutator phentoype tumors. Related to **Figure 2**. **Table S2.** Gold standard datasets. Related to **Figure 2**.

**Table S3.** Gold standard enrichment analysis with OncoMerge predicted somatic mutations. Related to **Figure 2**.

**Table S4.** Gold standard enrichment analysis with OncoMerge predicted somatic mutations using randomized somatic mutation matrices to show that overlap is not due only to seed genes. Related to **Figure 2**.

**Table S5.** Relative mean contribution of each mutation type to final mutation frequency. Related to **Figure 2**.

**Table S6.** Gold standard enrichment analysis from OncoMerge sensitivity analyses for cutoffs and parameters. Related to **Figure 3**.

**Table S7.** Mechanistic and causal regulatory interactions from the TCGA Pan-Cancer SYGNAL network. Related to **Figure 5**.

**Table S8.** Active TF regulatory network from TCGA PanCancer SYGNAL network. Related to Figure 6.

**Table S9.** Active TF regulatory network motifs. Related to **Figure 6**.

## Supplementary data

**Data S1.** OncoMerge summaries for all cancers. Related to **Figure 3**.

## Acknowledgments

This work was supported by NIH-NINDS Award # 1R01NS123038-01, and 1R01NS119650-01. The authors also acknowledge Robert Schultz for assistance in preliminary studies, and the Cancer Genome Atlas Research Network for the TCGA Pan-Cancer Atlas multi-omic patient tumor profiles.

## Author contributions

Conceptualization, C.L.P.; Methodology, S.S.S, S.F.W., and C.L.P.; Investigation, S.S.S and C.L.P; Visualization, S.S.S, S.F.W, E.M.L., S.A.O., and C.L.P.; Writing – Original Draft, C.L.P.; Writing – Review & Editing, S.S.S, S.F.W, E.M.L., S.A.O., and C.L.P.; Funding Acquisition, C.L.P.; Resources, C.L.P.; Supervision, C.L.P.

