## Supplementary Information for "Systematic integration of protein affecting mutations, gene fusions, and copy number alterations into a comprehensive somatic mutational profile"

### Supplementary figures

**Figure S1**. OncoMerge flow-chart that describes how the putative protein affecting mutation (PAM), transcript fusions (Fusion), and putative copy number alteration (CNA) data are integrated and filtered to generate an integrated mutation matrix. Related to **Figure 1**.

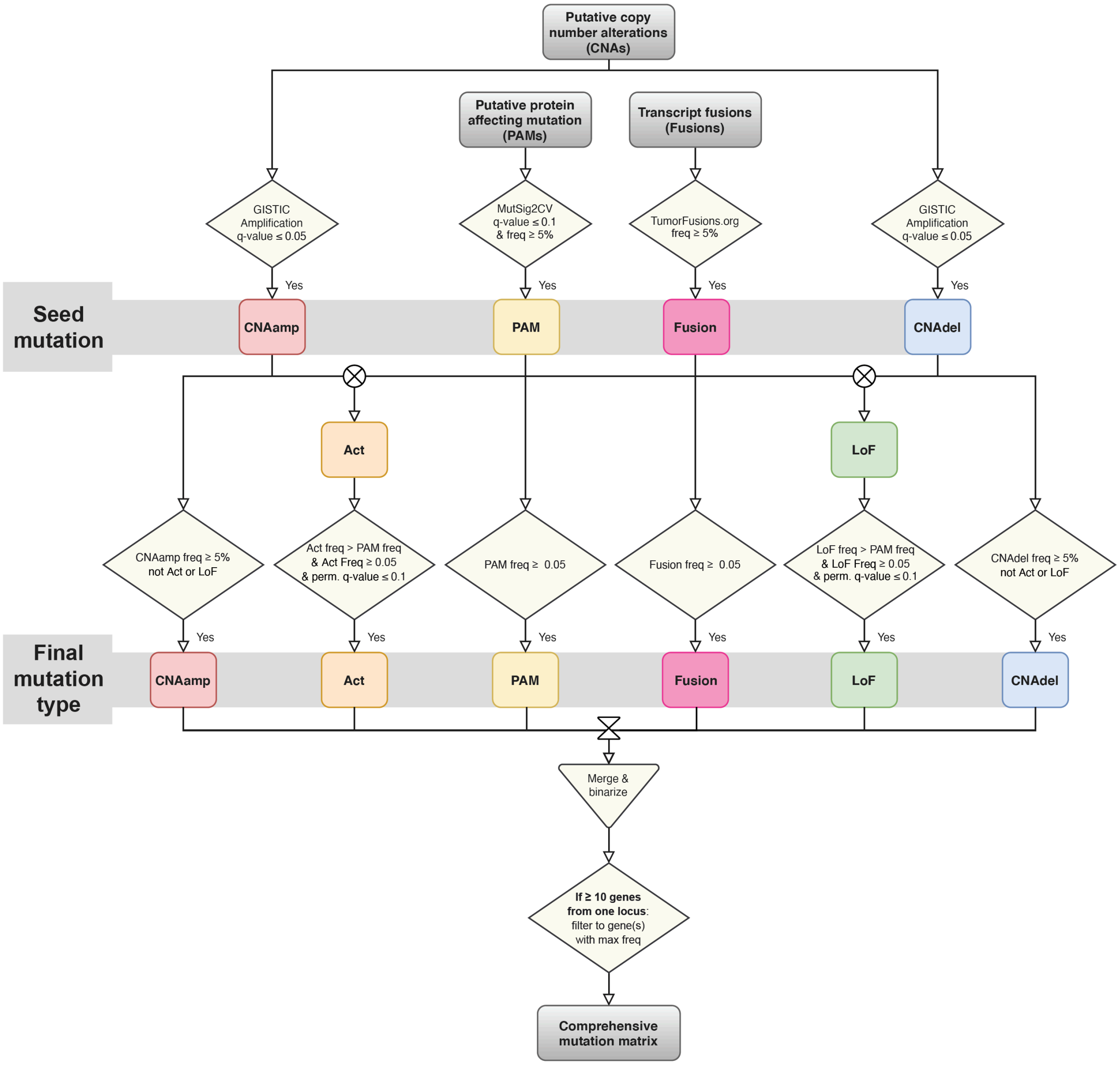

**Figure S2.** Average contribution of somatic mutation type to the final mutation frequency. Related to **Figure 2**.

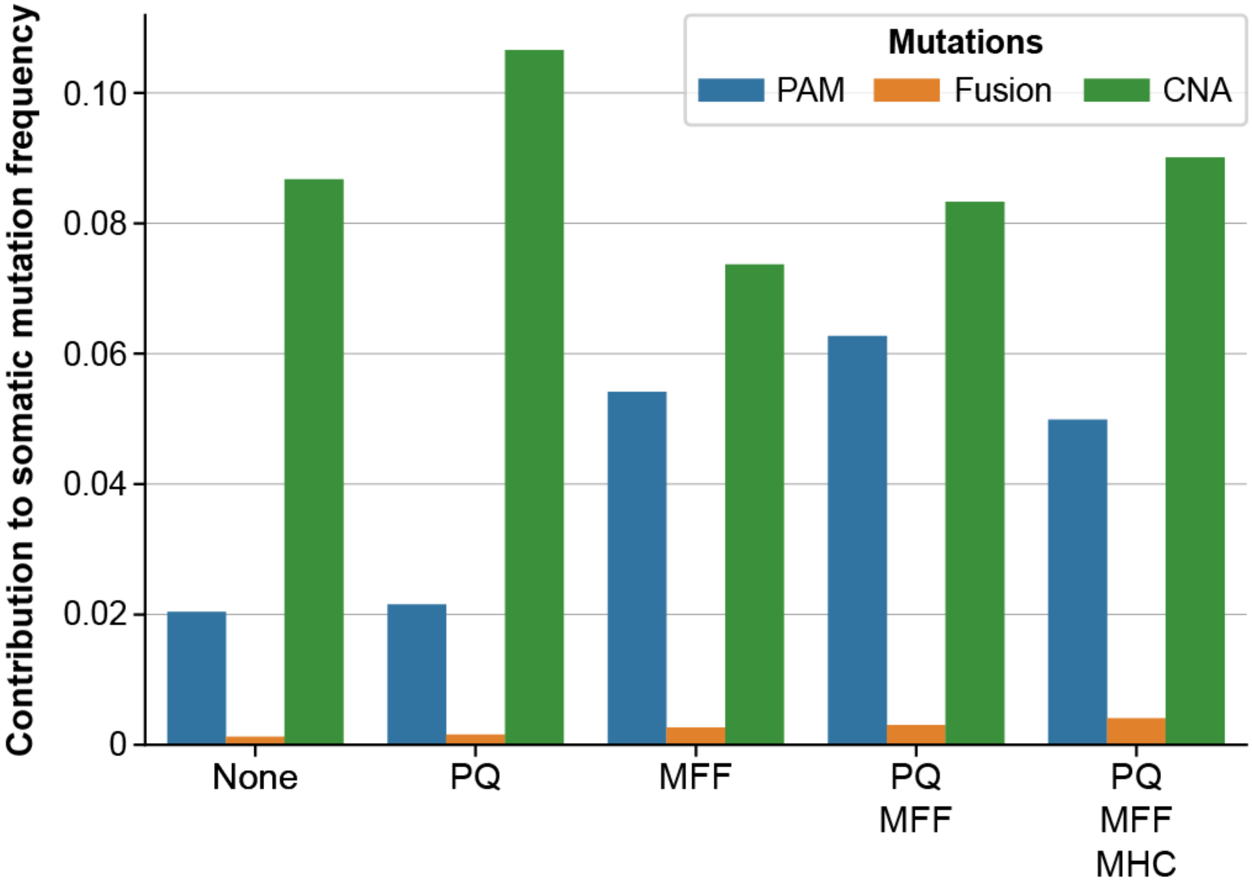
